# Strain-dependent assessment of Powassan virus transmission to deer ticks

**DOI:** 10.1101/2023.07.13.548769

**Authors:** Rebekah J. McMinn, Samantha Courtney, Sam R. Telford, Gregory D. Ebel

## Abstract

Powassan virus (POWV) is an emergent tick-borne encephalitis virus of Lyme disease endemic sites in North America. Due to range expansion and local intensification of deer tick vector (*Ixodes scapularis*) populations in the northeastern and upper midwestern U.S., encephalitis cases are increasingly being reported. A better understanding of the transmission cycle of POWV may allow for predicting the eventual public health burden. Recent phylogeographic analyses of POWV have identified geographical structuring, with well-defined northeastern and midwestern clades of the deer tick virus subtype (lineage II); sublineages exist within each clade. It may be that the local sublineages differ in their capacity to be transmitted by the deer tick vector. Accordingly, we determined whether there are strain-dependent differences in transmission. Five recent, low passage POWV isolates were used to measure aspects of vector competence, using viremic and artificial infection methods. Infection rates in experimental ticks remained consistent between all five isolates tested, resulting in 12-20% infection rate and no clear differences in viral load. We conclude that there is a genotype independent ability of POWV to infect deer ticks, and that differences in transmission efficiency are not likely to serve as the basis for regional differences in apparent public health burden.

## Introduction

Tick-borne diseases are of increasing concern due to the expanding range and density of ticks and the pathogens they carry^1–3^. In North America, one of these pathogens is Powassan virus (POWV; *Flaviviridae*), which can cause severe neuroinvasive disease in humans similar to its Eurasian relative, tick-borne encephalitis virus (TBEV). In the United States, POWV human cases have increased from 0.9 cases per year (1958-2007) to 16.7 cases per year (2008-2021)^4,5^.

POWV (lineage I) was originally identified in enzootic tick vectors that infrequently bite humans and thus was a rare infection. A second lineage of the virus (deer tick virus; DTV) was isolated from deer ticks in the late 1990s, and has since been shown to be endemic in the Northeast and Midwest U.S.^6,7^. Natural infection of deer ticks has significant implications for human disease: this species is an aggressive-human biter and regarded as the most medically important tick vector in North America, transmitting four other zoonotic pathogens including *Borrelia burgdorferi*, the bacterial agent of Lyme disease^8^. Increasing POWV seropositivity in deer^9^, recent reports of high infection rates in ticks^10–12^, and human cases reported in states with no prior history of disease^5,11,13,14^ suggest the potentially dynamic epidemiology of this virus.

Phylogeographic analyses of POWV genomes suggest that the virus exists in discrete transmission foci with infrequent dispersal, as with TBEV in Eurasia. Such isolation could imply adaptation to local transmission conditions such as host or vector diversity and genetic background, as well as extrinsic influences (microclimate). Indeed, geographic risk of human POWV disease appears to be heterogeneous, with Massachusetts, Wisconsin, and Minnesota, Maine, and New York accounting for >90% of all cases. This contrasts with the known risk for Lyme disease, with intense zoonotic transmission throughout all the New England states southward to Pennsylvania and Maryland, as well as in Wisconsin and Minnesota. It may be that there is differential risk associated with local viral genotypes; this is evident for TBEV in Western Europe. Accordingly, to determine whether local POWV strains may differ in their capacity to be transmitted, we measured the capacity for five isolates of POWV to infect deer ticks.

Experimental studies of POWV in ticks have been performed in *Dermacentor variabilis*^15,16^, *Haemaphysalis longicornis*^17^, *Amblyomma americanum*^16^, and *I. scapularis*^16,18^. Notably, these studies have relied on the use of highly passaged historical virus strains. With the emergence of POWV in North America, the use of low-passage, contemporary, genetically, and geographically diverse isolates is essential for an accurate representation of tick transmission.

## Materials and Methods

### Viruses and plaque assays

BHK-21 (ATCC CCL-10) cells were grown in DMEM supplemented with 10% FBS and 1% Penicillin/Streptomycin. L929 cells were grown in DMEM supplemented with 5% FBS and 1% Penicillin/Streptomycin. POWV isolates were grown up in BHK cells for 3-6 days and tittered via plaque assay on BHK cells.

### Mice and ticks

BALB/c (strain #000651) mice were obtained from Jackson Laboratories. Approval for animal protocols was obtained by the Colorado State University Institutional Animal Care and Use Committee, protocol 1257. Ticks were obtained from the Oklahoma State University tick rearing facility and kept at 24°C with 90-95% RH in glass humid chambers with ∼ 2 inches of saturated potassium sulfate. Ticks were kept in 5-dram polystyrene vials with mesh sieve tops. Larval ticks were obtained by allowing replete adult female *I. scapularis* to oviposit. Eggs were separated into ∼200 egg bunches in separate vials and allowed to hatch. Ticks were washed in 70% EtOH and PBS and moved into a clean vial and humid chamber every 3-5 weeks.

### Nucleic acid extraction and qRT-PCR

50μL sample was used for nucleic acid extraction using the Omega viral RNA/DNA kit on a King Fisher robotics platform. qRT-PCR was performed using the EXPRESS One-Step SYBR GreenER kit (Invitrogen) following manufacturer’s instructions with 2μL sample volume with the following primers directed towards the NS5 gene: fw: GGCCATGACAGACACAACAGCGTT TG and POWV rv:GAGCGCTCTTCATCCACCAGGTTCC. Melt curves were used to confirm true POWV positives over background.

### Viremic transmission

Larval ticks (separated into vials with ∼200 larvae) were used to infest 15-week-old female BALB/C mice. During infestation, mice were restrained in plastic restrainers and their tails were taped to prevent them from escaping. A single vial of larval *I. dammini* were added to the nape and upper back of each mouse. The infested mouse was kept in the restrainer, loosely wrapped in paper towels, and placed in secondary containment for 30 minutes. Mice were housed individually in wire-bottom cages with ∼1/2 inch of water placed in the bottom during the study. The next day (day 1), mice were inoculated intraperitonially with 1×10^4^ or 1×10^3^ PFU virus. Cage water was changed daily. Blood was collected via retro-orbital bleed on day 4 post infestation. As larval ticks became replete, they were collected from the water, surface sterilized in 70% EtOH and PBS and homogenized immediately or moved into 5-dram vials to be held as described above until they molted to nymphs. Eight weeks post repletion, molted nymphs were bead homogenized in tick diluent (200μL PBS supplemented with 20% FBS, 1% Penicillin/streptomycin, 2.5ug/mL amphotericin B, 50μg/mL gentamicin) at 24Hz for 2 minutes. POWV infection was determined by qRT-PCR and plaque assay as described above.

### Mouse viremia time course

15-week-old BALB/C female mice were inoculated intraperitoneally with 1×10^3^ PFU. Three mice were euthanized for each isolate at days 1, 2, and 4. Brain, spleen and blood were collected. Brain and spleen tissue were homogenized in 10% weight/volume of tick diluent at 30Hz for 2 minutes. RNA extraction and qRT-PCR was performed as described above. DTV infection was determined through qRT-PCR and plaque assay. PFU equivalents were determined through serial dilution of stock RNA via qPCR. This study was repeated with 6 mice euthanized per isolate on days 1 and 2, and 3 mice per isolate on day 4. The data presented here is combined from both studies.

### Immersion infection method

Nymphal *I. scapularis* ticks were dehydrated overnight at 26°C at ∼45% RH. Nymphs were immersed the next day (day -1) in 4mL of 1×10^3^ PFU/mL POWV for 1 hour at 37° C with light vortexing every ten minutes. Ticks were washed twice in PBS and moved into a ventilated tube to dry overnight. Foam capsules were made as previously described^19^ with 20mm outer diameter and 12.5mm inner diameter. The backs of 10-week-old male BALB/C mice were closely shaved and wiped with 70% EtOH. Two foam capsules were fixed the day before tick infestation (day -1) to each mouse with non-toxic leather adhesive (Tear Mender). One capsule was placed on the nape and the second was placed immediately posterior near the middle back. Capsule attachment was checked daily and patched with glue if needed. The next day (day 0), 15-20 immersed nymphs were added to capsules. Infested nymphs were checked daily and removed from the capsule when replete by cutting open the plastic capsule top and re-sealing with plastic stickers as previously described^19^. Nymphs were left to molt in humid chambers as previously described. Two weeks after the start of molting, nymphs were homogenized as described above and screened via qRT-PCR and plaque assay.

## Results

### POWV isolates used in this study

Five POWV isolates were chosen based on geographic and temporal origin and passage history (Table 1). This includes a single lineage I isolate (M11665) and four lineage II isolates: two in the Northeast clade (NFS9601 and ME19-1051) and two in the Midwest clade (FA5/12-40 and NJ19-56). M11665 (lineage I; 1965) has been passaged moderately in cell culture and suckling mice. Several isolates (ME19-1051, NJ19-56, and FA5/12-40) have been passaged no more than twice in BHK cells. Notably, NJ19-56 was collected in New Jersey, though clusters genetically with the Midwest isolates as discussed previously^20^.

**Table 1.** Powassan virus isolates used in this study. SM= suckling mice; V=Vero cells; B=BHK cells; P=unknown passage; lineage I = POWV; lineage II = DTV

| Isolate ID | Lineage | location | year | source | passage history | GenBank accession |
| --- | --- | --- | --- | --- | --- | --- |
| M11665 | I | Ontario | 1965 | *I. cookei* | P1SM1V1B1 | OP823404 |
| NFS9601 | II | Nantucket, MA | 1996 | *I. scapularis* | SM1B2 | HM440559 |
| ME19-1051 | II | Rockland, ME | 2019 | *I. scapularis* | B1 | OP823442 |
| FA5/12-40 | II | Spooner, WI | 2008 | *I. scapularis* | B2 | OP823475 |
| NJ19-56 | II | Hardwick, NJ | 2019 | *I. scapularis* | B1 | OP823460 |

### Viremic transmission to ticks

Viremic transmission of POWV was assessed in *I. dammini* ticks fed on BALB/c mice infected intraperitoneally with 10^3^ PFU (Fig. 1A). Approximately 200 larvae were infested a day prior to infection (day 0) to better overlap peak viremia with blood-feeding. Three mice were infected per isolate on day one and replete larvae were collected as they detached, largely on days three and four. Ten or approximately 20% of replete larvae per mouse were immediately homogenized and used to determine rates of virus acquisition. 90.0% of ticks had detectable levels of viral RNA, and10.0% had detectable virus. Remaining larvae were left to molt for eight weeks and nymphs were homogenized and screened for POWV. A median of 35 (range 10-78) nymphs were obtained per mouse. Blood was collected from all mice on day four, though viral RNA was not detected via qRT-PCR.

**Figure 1.**
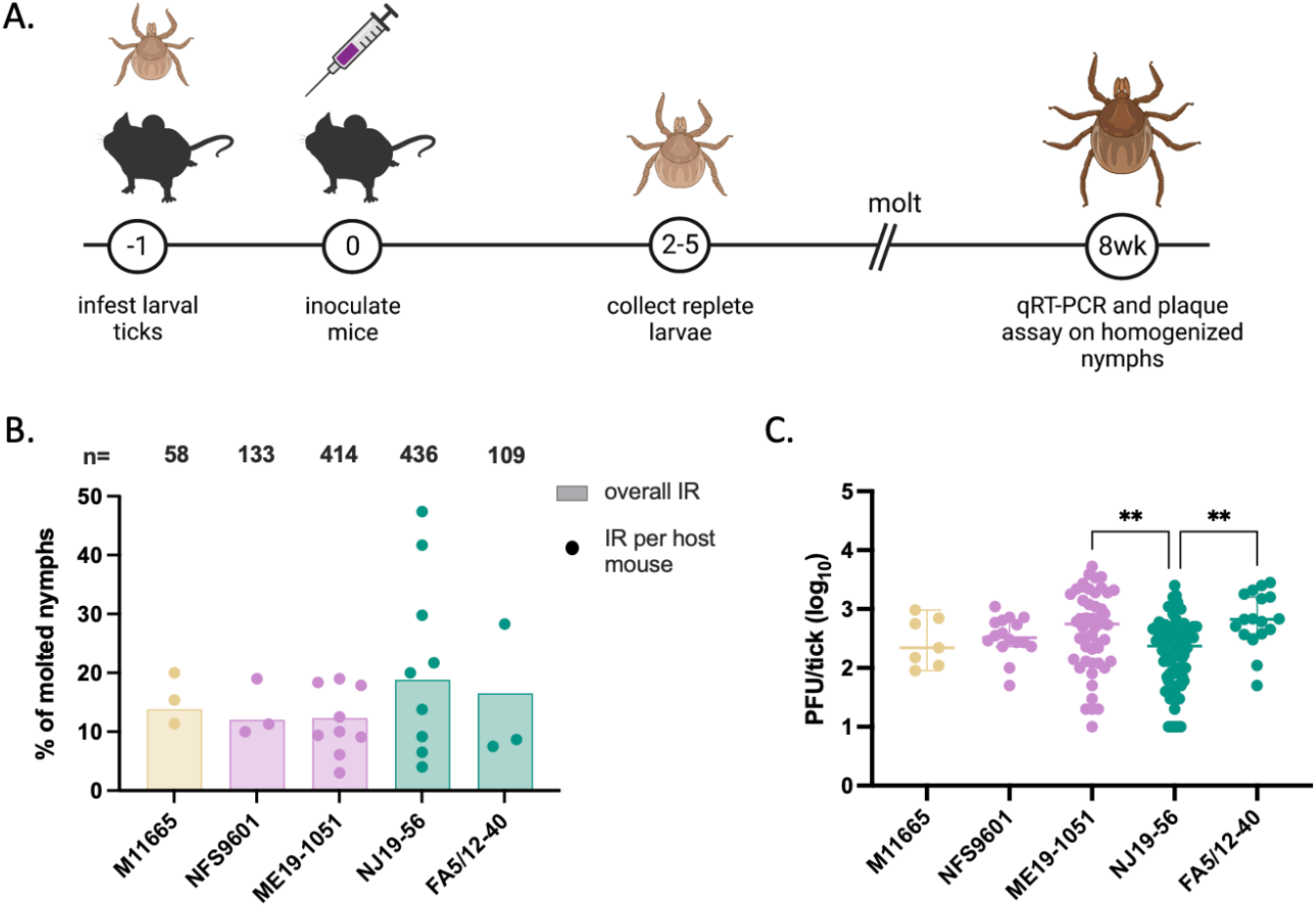
Viremic transmission to ticks. A) Methodology of viremic transmission experiments. B) Nymph infection rates per isolate. Colored bar represents overall IR, colored circles represent IR for nymphs fed on the same host mouse. Total n is listed at the top. Kruskal-Wallis test p=0.55 C) Viral load of all nymphs Kruskal-Wallis test **p<= 0.001

To bolster our sample size, the experiment was repeated at a lower inoculum (due to viral titers of our stocks) using only two isolates: ME19-1051 and NJ19-56 and six replicate mice per isolate. All replete larvae were allowed to molt and harden and were analyzed 8 weeks after the larval bloodmeal. A median of 54.5 (22-90) molted nymphs were obtained from each mouse. No statistical significance in viral titer or infection rate (IR) in nymphs was observed between studies (Table 2) and results from the combined experiments are discussed below.

**Table 2.** Statistical analysis between first and second viremic transmission experiments.

|  |  | exp. 1 | exp. 2 | test | p value |
| --- | --- | --- | --- | --- | --- |
| ME19 #1051 | median PFU | 370 | 610 | Mann-Whitney t-test | 0.1166 |
|  | mean proportion | 11.33 | 11.9 | t-test with Welch's correction | 0.9007 |
| NJ19 #56 | median PFU | 250 | 220 | Mann-Whitney t-test | 0.9243 |
|  | mean proportion | 36.93 | 13.88 | t-test with Welch's correction | 0.0758 |

No significant difference in nymph IR was determined between POWV isolates (Kruskal-Wallis test). The overall IR averaged 14.7% (range 12.0-18.8) amongst all isolates. Greater IR variation was observed between ticks collected from individual host mice. For NJ19-56, tick IRs between host mice ranged from 4.0 to 47.4% (Fig. 1B). Some subtle trends in nymph viral loads were observed between isolates. M11665, NFS001, and NJ19-56 resulted in an average 4×10^2^ PFU/tick, while ME19-1051 and FA5/12-40 average 1×10^3^ PFU/tick, though this difference is only significant compared to NJ19-56 (Kruskal-Wallis test with Dunn’s multiple comparisons; Fig. 1C).

### Viremia in BALB/c mice

To assess differences in the ability to establish viremia in the vertebrate host, female BALB/C mice were infected with 10^3^ PFU and three mice per isolate were euthanized one, two, and four days post infection (dpi) (Fig. 2). Due to limits of virus detection in the blood, PFU equivalents were calculated by extrapolation from qRT-PCR standard curves. Viral titers in the blood were highest one dpi averaging 9×10^2^ PFU eq/mL, dropping roughly ten-fold two dpi and only detected at low levels four dpi. Viral RNA was detected in the spleens of all mice at all time points, highlighting successful infection. In the brain, viral RNA is detected by four dpi, and rarely at early time points of infection for NJ19-56. In the blood at one and two dpi, 83.3% and 94.4% of mice respectively had detectable levels of POWV RNA. Two dpi, significantly higher PFUeq/mL (was determined in the blood of ME19-1051-infected mice. By four dpi, viral loads in the blood drop off: 50.0% of mice are determined POWV RNA positive with very low approximate viral loads less than 10 PFUeq/mL.

**Figure 2.**
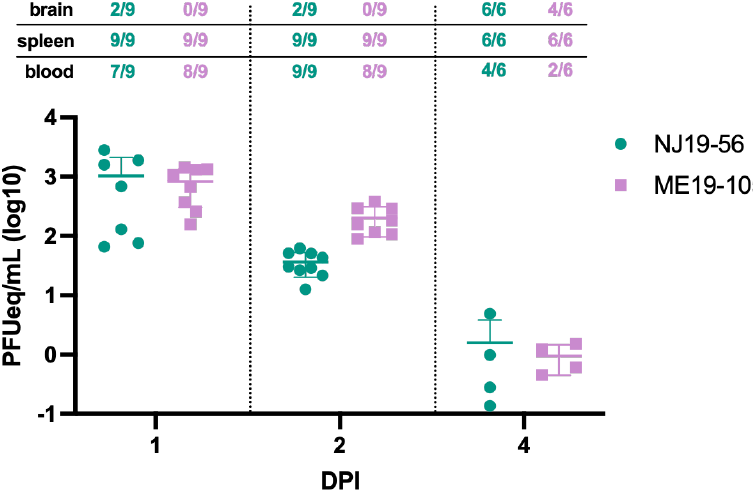
Viremia of DTV in BALB/c mice. Blood, brain, and spleen was taken from mice euthanized on days 1 (n=9), 2 (n=9), and 4 (n=6) post infection. Proportions of POWV RNA positive tissues are indicated in the table above the graph.

### Artificial infection of ticks

To determine differences in the ability of isolates to infect the tick without host-dependent viremia, artificial infection using two DTV isolates (NJ19-56 and ME19-1051) was used. *I. scapularis* nymphs were immersed in 10^3^ PFU/mL of DTV, fed on naive mice, and allowed to molt to adults for eight weeks before processing (Fig. 3A). A subset of immersed, unfed nymphs were screened for DTV RNA by qRT-PCR four weeks post-immersion, though only one tick (2.6%) was found positive. Artificial infection of blood-fed nymphs resulted in 15.0-20.5% IR in adults with viral loads averaging 7.5×10^3^ PFU (Fig. 3B and C). Differences between isolates (viral load and IR) were not significant (Kolmogorov-Smirnov test and chi square test respectively) and no differences were determined between male and female infected ticks.

**Figure 3.**
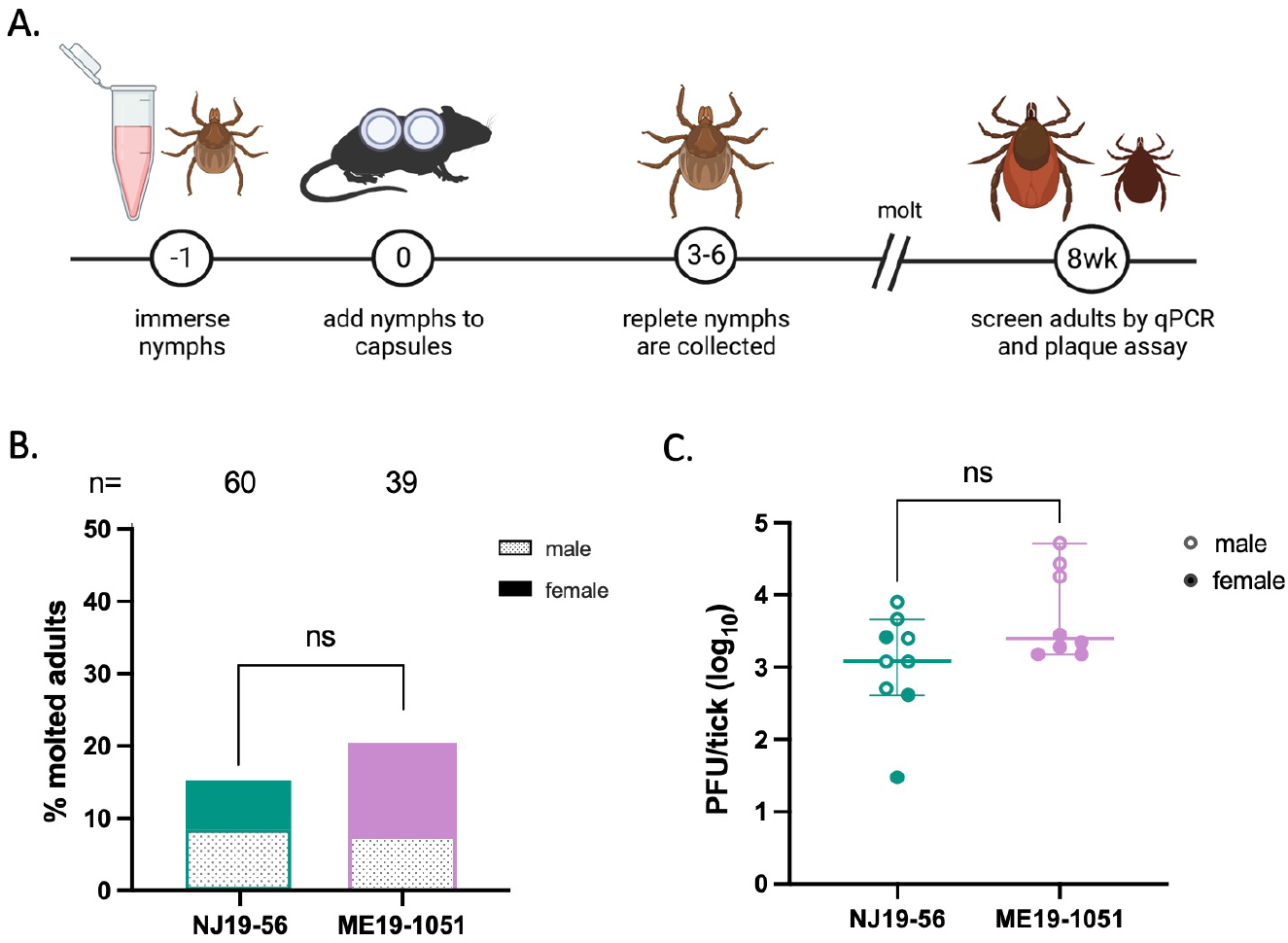
Artificial infection of ticks. A) Methodology of artificial infection. B) Adult male and female infection rates per isolate. Total n is listed at the top. Chi square test p= 0.48 C) Viral load of all nymphs Kolmogorov-Smirinov test p=0.56

## Discussion

We sought to determine whether POWV strains differed in their capacity to transmit to and infect ticks. Using five low-passage, genetically diverse POWV isolates of different geographic and temporal origin, we compared the rate of viremic transmission and infection success in a single *I. scapularis* reference strain, thus holding the tick genetic background constant. Comparable infection rates were observed between all isolates regardless of infection method, suggesting viral mechanisms of survival and transmission within ticks may be genotype independent.

Ticks were infected via two methods: viremic transmission (feeding on an infected mouse) and artificial infection (via immersion). During viremic transmission, the virus establishes infection in the blood of the host animal and is ingested into the midgut of the engorging tick.

Artificial infection removes the variable factor of host viremia; however, blood-feeding is still necessary to induce physiological changes in the tick that promote virus infection^21,22^. This effect was observed in our data as only 2.6% of un-fed nymphs were DTV-positive, compared to 17.8% of blood-fed molted nymphs. Therefore, the combination of viremic and artificial infection allows us to determine the impact of host viremia on IR and assess strain-dependent differences in the ability to infect ticks. Surprisingly, both methods resulted in comparable overall tick IRs between 10-20%, similar to previously published data^16,18^; however tick IR was variable between individual host mice. Artificial infection resulted in somewhat higher viral loads per tick, though this did not seem to increase the likelihood of infection. Given these results, it seems infection acquired by blood feeding on a live animal can result in variable IR depending on the individual animal, but that overall IR is similar to artificial feeding when results from several animals are aggregated.

Despite significant genetic and ecological variability, the overall tick IR was also similar between isolates, regardless of infection method. This is surprising given the number of barriers a virus must overcome to (1) develop sufficient viremia in the host mouse, (2) establish infection in the tick, and (3) survive the histolysis and rearrangement of tissues that occurs during molting^23–25^. With mosquito-transmitted viruses, a threshold viremia determines vector infection; although we did not define such a threshold, all strains exceeded a putative minimum viremia threshold for tick infection. Some differences in host viremia were observed between two DTV isolates: ME19-1051 and NJ19-56, with higher approximate viral loads observed two dpi for ME19-1051-infected mice. Though this difference did not seem to contribute significantly to tick infection rates, significantly higher viral loads were observed for ME19-1051-infected ticks and this trend was repeated via artificial infection. Such differences can impact viral maintenance or subsequent transmission and may have implications for human disease as inoculum size may modulate pathology as observed with other arboviruses. Finally, all isolates were successfully transstadially transmitted (TST). Though only a small number of larvae were screened immediately post-repletion, the proportion that were virus-positive is similar to the IR in nymphs post-molt, indicating low barriers to TST as previously described^18,26^. Together, these results suggest that mechanisms of POWV infection in *I. scapularis* are not POWV genotype dependent and may be highly conserved.

We also assessed the infection rate and viral load of male and female ticks via artificial infection. Previously, our group reported significantly more female DTV-positive ticks collected form the Northeast U.S.^20^. Though we did not observe differences in the present data, our sample size is relatively small and could be impacted by the method of infection. Biological factors, such as the difference in sex organs and the size and function of the midgut and salivary glands^27^ could influence virus replication and survival. Thus, additional work is needed to determine virus tropism in adult ticks.

In conclusion, we determined consistent rates of infected ticks between diverse viral isolates via viremic and artificial infection. The minimal strain-dependent differences in tick transmission that we observe for POWV suggests that such variation is not the basis for differential human risk (i.e., sites where POWV is expected based on the enzootic presence of the virus and intense Lyme disease risk, but human cases rarely reported). Variation in enzootic viral transmission may be more closely associated with variables related to the vertebrate host.

## Notes

### Competing Interest Statement

The authors have declared no competing interest.

